# Scheduling problems and the energetics of biparental care in a model of imperiled seabirds

**DOI:** 10.64898/2026.08.08.743669

**Authors:** Liam U. Taylor, Patricia L. Jones, Mark F. Haussmann, Robert A. Mauck

## Abstract

For organisms with biparental care, successful reproduction hinges on coordination between partners. Seabirds face an extreme coordination challenge because parents must schedule nest attendance on land with long-distance foraging trips at sea. We present a computational model of incubation schedules for a vulnerable seabird, the Leach’s Storm-Petrel (*Hydrobates leucorhous*). Using only simple energetic rules and parameters, the model recapitulates natural incubation rhythms, exposes a tradeoff between parent energy and egg attendance, and predicts severe reproductive failure in harsh environments. Incubation primarily fails through “schedule breakdown” — a single point in the season when both parents spend too long foraging and the egg dies from cold. The resilience of the developing offspring to neglect is thus a fundamental adaptation to the uncertainties of biparental care. These results raise new alarms about the indirect causes of reproductive failure in sensitive marine species and provide theoretical foundations for the evolutionary ecology of scheduling behaviors.

## INTRODUCTION

Parents make decisions about how much time and energy to invest in their offspring rather than save for themselves (Clutton-Brock 1991; Stearns 1992). For organisms with biparental care, reproductive success depends on the energetic decisions of both parents (Royle *et al*. 2016; Trivers 1972). Although some biparental care systems involve two parents that each specialize in different forms of care (e.g., one parent attends the nest while the other defends the territory, as in some cichlids; Itzkowitz *et al*. 2001), others require parents to coordinate in a single form of care (e.g., switching off attending the nest, as in most shorebirds; Székely & Reynolds 1995). The result is a rhythm, routine, or schedule of alternating reproductive investment played out over the course of a breeding season (Dominoni *et al*. 2017; Houston & McNamara 1999).

Empirical studies document biparental care schedules across a wide range of organisms (e.g., birds: Bulla *et al*. 2016; frogs: Schulte & Summers 2022; primates: Wood & Fernandez-Duque 2026), while studies and experiments with game theory provide insights into how parents should make optimized decisions in response to partner behaviors (e.g., Dearborn 2001; Johnstone *et al*. 2014; McNamara *et al*. 1999). However, there is a striking diversity in biparental care schedules among even closely related taxa (Bulla *et al*. 2016). We lack a first-principles approach to understanding how or why particular schedules emerge, operate, or break down for individuals, pairs, or species.

Seabirds face an extreme scheduling challenge, because their foraging grounds are far from their nests. Consequently, all seabirds exhibit biparental care (Cockburn 2006; Lack 1968). While one parent forages at sea, the other can attend the nest. Well-coordinated pairs establish consistent schedules that keep eggs warm or keep chicks fed (e.g., Chaurand & Weimerskirch 1994; Davis 1982; Tyson *et al*. 2017). Yet seabird parents must contend with both limited information about their traveling partners and the fundamental uncertainty of marine environments. As these environments become less productive and predictable, disruptions to breeding success and long-term partnerships raise serious concerns for seabirds (Sun *et al*. 2024; Ventura *et al*. 2021) at a time when nearly all seabird populations are already in decline (Dias *et al*. 2019).

Here, we use a computational model of biparental care to investigate incubation scheduling for a vulnerable species, the Leach’s Storm-Petrel (*Hydrobates leucorhous*). This small, nocturnal seabird has particularly well-studied parental care patterns (e.g., Elliott *et al*. 2021; Haussmann *et al*. 2024; Tyson *et al*. 2022; Zangmeister *et al*. 2009). A breeding storm-petrel must coordinate with its partner for over a month to incubate an egg in an underground burrow while traveling hundreds of kilometers to forage at sea (Hedd *et al*. 2018). Our model simulates two parents as they follow energetic rules for switching between incubating and foraging. Parameterizing the model with previously published data from the Northwest Atlantic, we ask three questions: (1) are simple energetic rules sufficient to generate natural schedules, or are more complicated social dynamics required? (2) what energetic strategies allow two parents to successfully schedule care? and (3) why, and under what environmental conditions, do schedules fail?

## METHODS

### Study system

Leach’s Storm-Petrels (Procellariiformes: Hydrobatidae: *Hydrobates leucorhous*) are small (50 g), long-lived (30+ yr), sexually monomorphic seabirds (Pollet *et al*. 2021). Like other procellariiforms, such as albatrosses, petrels, and shearwaters, Leach’s Storm-Petrels are pelagic specialists with enduring pair-bonds and low extra-pair paternity (Mauck *et al*. 1995; Warham 1990). Populations nest on island colonies across the North Atlantic and North Pacific, migrating to southern oceans during the non-breeding season (Pollet *et al*. 2019). The nests are underground burrows with single-egg clutches. Parents forage far offshore for patches of small fish, crustaceans, and zooplankton near the surface of the water (Hedd *et al*. 2018; Watanuki 1985). The species is currently recognized as IUCN Vulnerable, with declining populations and conservation threats that include predation and disturbance at island colonies along with light pollution, oil spills, and prey depletion at sea (Pollet *et al*. 2023).

### Computational model

We developed an agent-based model of biparental care during the seabird incubation period (Fig. 1). The model simulates the daily energetic flux of two parents as they lose energy through metabolism and gain energy through foraging. The model is run in C++11, with analysis and visualization of simulation results in R 4.6.0 (R Core Team 2026; Wickham *et al*. 2019).

**Figure 1.**
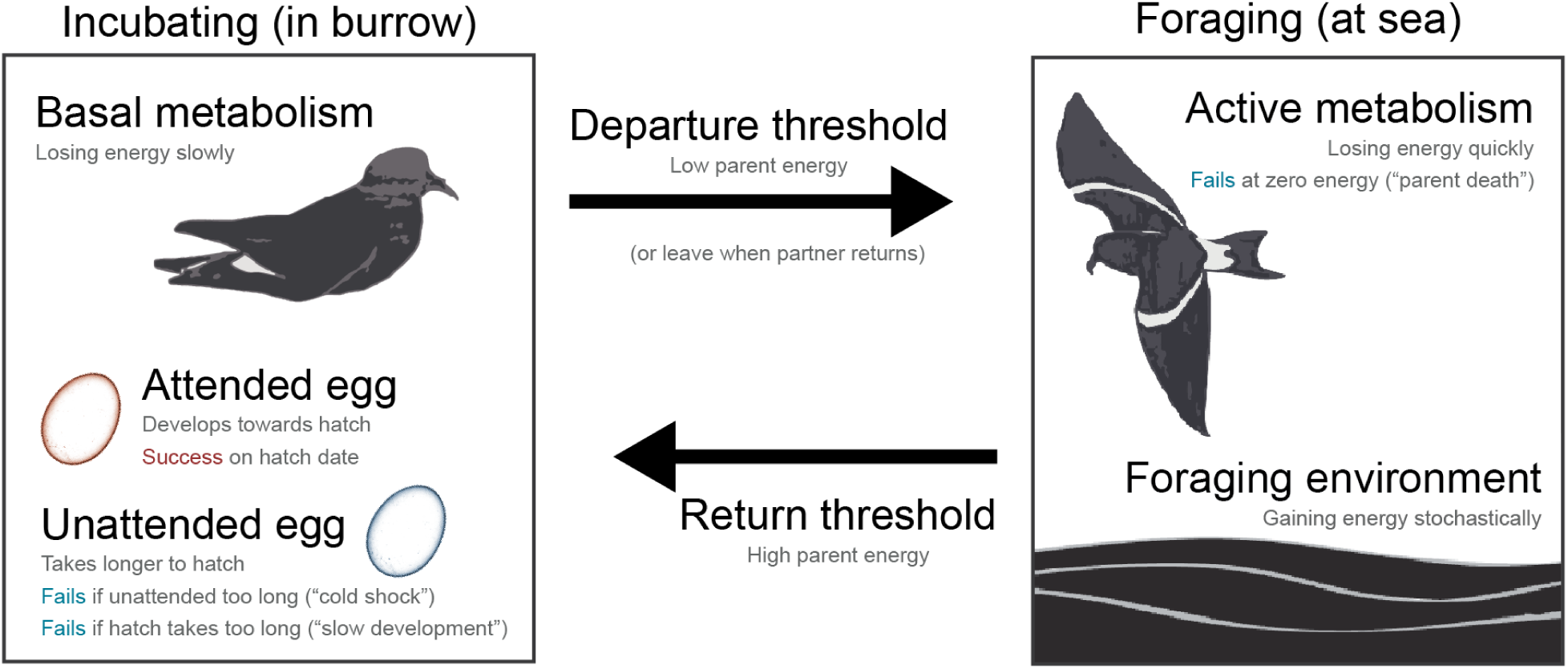
Model structure for a simulation of biparental incubation in Leach’s Storm-Petrels (*Hydrobates leucorhous*). Incubating parents lose a fixed amount of energy each day until reaching a low energy (departure) threshold, at which point they switch to foraging. An incubating parent departs immediately if their partner returns. Foraging parents lose energy more rapidly but draw energy from a normally distributed foraging environment, switching to incubation upon reaching a high energy (return) threshold. A foraging parent dies at zero energy. An attended egg is warm and develops towards hatching when an incubating parent is present. An unattended egg is cold, takes longer to hatch, and dies after too many consecutive days of neglect. Each incubation period ends in one of four outcomes: successful hatch, failure from an overlong incubation period (“slow development”), failure from too many consecutive days when the egg is unattended (“cold shock”), and failure from parent death.

Model and analysis code is available on GitHub (https://github.com/ltaylor2/Petrel_Schedule). The model simulates the daily activity of three agents (a female parent, a male parent, and an egg) during incubation. Each agent follows simple energetic rules. Parents can be in one of two states: *incubating* in the nest or *foraging* at sea. While *incubating*, a parent loses energy at a fixed, basal metabolic rate. While *foraging*, a parent loses an even greater amount of energy from active metabolism. However, a *foraging* parent can also gain energy from the environment, sampling once per day from a normal distribution with a characteristic mean and standard deviation, with negative draws clamped to zero. Stochasticity in the model emerges from this uncertainty in daily foraging outcomes. A parent dies if it has ≤0 energy at the end of the day.

At the end of each day, parents have the opportunity to switch between *incubating* and *foraging* based on energetic thresholds. When the energy of an *incubating* parent drops too low (the *departure* threshold), it switches to the *foraging* state before the start of the next day. Similarly, if the energy level of a *foraging* parent rises high enough (the *return* threshold), it switches to the *incubating* state before the next day. To account for travel time, parents must forage for at least two days before returning. Finally, only one parent incubates at a time. Thus, an *incubating* parent will switch to *foraging* if its partner returns to the nest, even if it has not yet reached its *departure* threshold. If both parents switch from *foraging* to *incubating* on the same day, one is chosen at random to return to *foraging*.

The egg sits in the nest, developing towards its *hatch date*. The egg can be in one of two states: *attended* or *unattended*. The egg is *attended* whenever a parent is *incubating*, which warms the egg and moves it one day closer to hatching. If neither parent is *incubating*, then the *unattended* egg is cold. Unattended, cold eggs have slower development, taking longer to hatch (*cold delay*) and failing altogether if they experience too many consecutive cold days (*cold tolerance*; Boersma & Wheelwright 1979; Elliott *et al*. 2021). Although the sensitivity of eggs to cold likely varies across development (Ahmad & Li 2023), this variation is not quantified in storm-petrels and we assumed that neglect has equal consequences across the incubation period.

Each simulation had four possible outcomes: (1) successful hatch, (2) failure due to “slow development” of the egg, (3) failure due to “cold shock” of the egg, and (4) failure due to parent death. The egg fails from “slow development” by experiencing too many cumulative days unattended (i.e., when *hatch date* is pushed beyond a *hatch date limit* via accumulated *cold delays*). Note that in this model, the only way that hatch date is extended is through a *cold delay*. The egg fails from “cold shock” when it experiences too many consecutive days unattended (i.e., pushed beyond the *cold tolerance*). Parent death is counted as another form of reproductive failure because both parents are generally needed to feed the chick after hatch. A simulation stopped immediately when the egg hatched, the incubation period hit the *hatch date limit*, the egg died, or a parent died, whichever occurred first.

### Biological parameters

We parameterized the model using energetic values for populations of Leach’s Storm-Petrels in the Northwest Atlantic (Table S1). We estimated several egg parameters using unpublished data from a long-term study of Leach’s Storm-Petrels on Kent Island, New Brunswick, Canada, or from the congeneric Fork-tailed Storm-Petrel (*Hydrobates furcatus*; Boersma & Wheelwright 1979; Table S1). For empirical parent strategies, we included a wide range of estimates around energy measured at the beginning of incubation bouts for *return* thresholds and at the end of incubation bouts for *departure* thresholds (Ricklefs *et al*. 1986). Both values were measured at the breeding ground and therefore account for the costs of travel. Given the lack of recent data on environmental conditions, we used older data from a Newfoundland population (Montevecchi *et al*. 1992). These historical environmental parameters may overestimate current conditions because several axes of marine productivity in the region are at risk of decline (Frumhoff *et al*. 2008; Pershing *et al*. 2021).

Leach’s Storm-Petrels are sexually monomorphic except for minor and overlapping variation in morphology (Pollet *et al*. 2021). However, there is evidence for subtle sexual differences in behavior, including increased egg attendance and shorter foraging trips by males (Mauck *et al*. 2011, 2023). Our model included only two fixed differences between females and males. First, the female parent pays the energetic cost of the egg at a single point at the beginning of the incubation period (Table S1). Second, because the female must lay the egg, she begins in the *incubating* state while the male begins in the *foraging* state.

### Model testing

First, we asked whether our model, which includes only simple energetic rules rather than complex or dynamic social strategies, can effectively represent the real incubation schedules of Leach’s Storm-Petrels. We compared model output to reported values from wild populations in the Northwest Atlantic: (1) hatching success rate, (2) average incubation bout length, (3) egg unattended rate, (4) hatch date, and (5) sexual bias in parental effort, as measured by the difference in average incubation bout lengths for males versus females (Table 1). The latter four metrics were calculated from successful incubation periods only. Incubation bout lengths excluded the first and last bouts for each parent, because the length of the first bout is sensitive to initial model conditions while the last bout is cut off at the hatch date. These comparisons used empirical parameters for the egg, environment, and the range of parent strategies. For all subsequent comparisons, parameters are kept at their empirical values unless otherwise noted.

**Table 1.** Model fidelity.

| **Result** | **Model estimate<sup>A</sup>** | **Reported value<sup>B</sup>** |
| --- | --- | --- |
| Hatching success | 90.4 ± 5.0% | 67.6–91.0%<sup>C</sup> |
| Incubation bout length<sup>D</sup> | 4.0 ± 1.3 days | 2.5–3.5 days<sup>C,E</sup> |
| Egg unattended rate<sup>D</sup> | 12.1 ± 2.6% | 7.3%<sup>F</sup> |
| Hatch date<sup>D</sup> | 46.0 ± 2.1 days | 42.6–45.3 days<sup>C,E</sup> |
| Male bias in incubation<sup>D</sup> | 0.2 days | 0.5 days<sup>G</sup> |

<sup>A</sup>Mean ± SD given empirical parameters for egg, environment, and the range of parent strategies;
<sup>B</sup>Range of mean empirical values available from Northwest Atlantic populations; <sup>C</sup>Pollet *et al.* (2021); <sup>D</sup>Estimates from successful incubation only; <sup>E</sup>Zangmeister *et al.* (2009); <sup>F</sup>Elliott *et al.* (2021); <sup>G</sup>Mauck *et al.* (2011).

Second, we used the model to investigate how different energetic strategies allow two parents to successfully schedule incubation. A “parent strategy” was defined as a unique combination of *departure* and *return* thresholds, which dictate how parents switch between states (Fig. 1). We tested parent strategies across combinations of *departure* thresholds (200–1,100 kJ, increments of 100 kJ) and *return* thresholds (400–1,200 kJ, increments of 100 kJ; Table S1). Excluding strategies where the *departure* threshold was greater than or equal to the *return* threshold, there were 54 individual parent strategies giving 2,916 pair strategy combinations.

Third, to investigate how incubation schedules and outcomes change based on environmental conditions, we reran simulations across a range of mean daily foraging intake values (130–170 kJ/day, increments of 10 kJ/day). As described below, these results led us to test the impact of egg cold tolerance on hatching success; we varied that parameter from 1 to 7 days in increments of 1 day and quantified success across the range of foraging conditions.

For each set of parameters, we ran 1,000 simulations and calculated the rate of four outcomes: success, failure from slow development, failure from cold shock, and failure from parent death. We also calculated two metrics that indicate potential fitness consequences extending beyond incubation. The first metric was mean parent energy (for successful incubation only), arbitrarily using the female as the focal parent to monitor for each pair. Parents may maximize lifetime reproductive success by maintaining their own condition, which allows them to care for chicks or survive to future breeding seasons (Stearns 1992). All else being equal, we thus expect that parents who maintain high energy levels will have higher lifetime reproductive success. The second metric was hatch date (for successful incubation only). Because cold storm-petrel eggs develop more slowly towards hatching (Boersma & Wheelwright 1979; Elliott *et al*. 2021) and seabirds that breed later in the season may experience lower chick condition and overall breeding success (Moreno 1998), we expect parents that hatch eggs more slowly will have lower lifetime reproductive success.

## RESULTS

### Model fidelity

The model effectively represented incubation behaviors in Leach’s Storm-Petrels (Table 1). Given empirical parameters for the environment, egg, and the range of parent strategies, hatching success averaged 90.4 ± 5.0%, which fell towards the top of the reported averages from wild populations (67.6–91.0%). Incubation rhythms were also generally accurate (see Fig. S1 for schedule visualizations). Although the average incubation bout length of 4.0 ± 1.3 days (mean ± SD) was at least 0.5 days longer than empirical averages (2.5–3.5 days), this species has incubation bouts that last up to six consecutive days (Pollet *et al*. 2021). Small changes to parameters could produce even more accurate incubation rhythms: for example, increasing the season starting energy by ∼18% (from 766 kJ to 900 kJ) reduced average incubation bout length to 3.4 days (Fig. S2).

The model overestimated egg neglect during incubation, with eggs unattended on 12.1% of days during successful incubation (Table 1). Although reports of early-season egg neglect in wild populations reach as high as 22% (Pollet *et al*. 2021), closely monitored pairs during successful incubation show an average of 7.3% (Elliott *et al*. 2021). Because unattended eggs develop more slowly, elevated rates of egg neglect in the model led to longer incubation periods (hatch date = 46.0 ± 2.1 days) than averages from wild populations (42.6–45.3 days).

Starting with only two fixed differences between sexes—the female pays the energetic cost of the egg, and the female begins the season incubating—the model also captured the subtle male bias in incubation effort reported for the species (Table 1). Averaged across the range of empirical parent strategies, male incubation bouts (4.1 ± 1.3 days) were slightly longer than those of females (3.9 ± 1.3 days), and male foraging bouts (5.1 ± 1.6 days) were slightly shorter than those of females (5.3 ± 1.6 days). Both fixed differences in the model contributed to this bias. Simulating a more expensive egg increased the contribution of males during incubation (Fig. S3). Flipping the starting states of males and females also moved the bias towards female incubation (Fig. S4).

### Parent strategies

Hatching success varied widely across parent strategies (Fig. 2A). Holding all other parameters at empirical values, hatching success rates for different pair strategy combinations ranged from 10.0% to 100.0% (5th–95th percentiles: 28.2–98.4%; Fig. S5). Parents with low departure or return thresholds were the most successful at hatching eggs (Fig. 2B). Parents with the lowest return thresholds were consistently successful, across the full range of corresponding departure thresholds or partner parameters, whereas the success of parents with low departure thresholds showed greater variation depending on the overall pair strategy combination (Fig. 2B). Empirical pair strategy combinations had high success compared to the full parameter search range but were not the most successful (Fig. S5).

**Figure 2.**
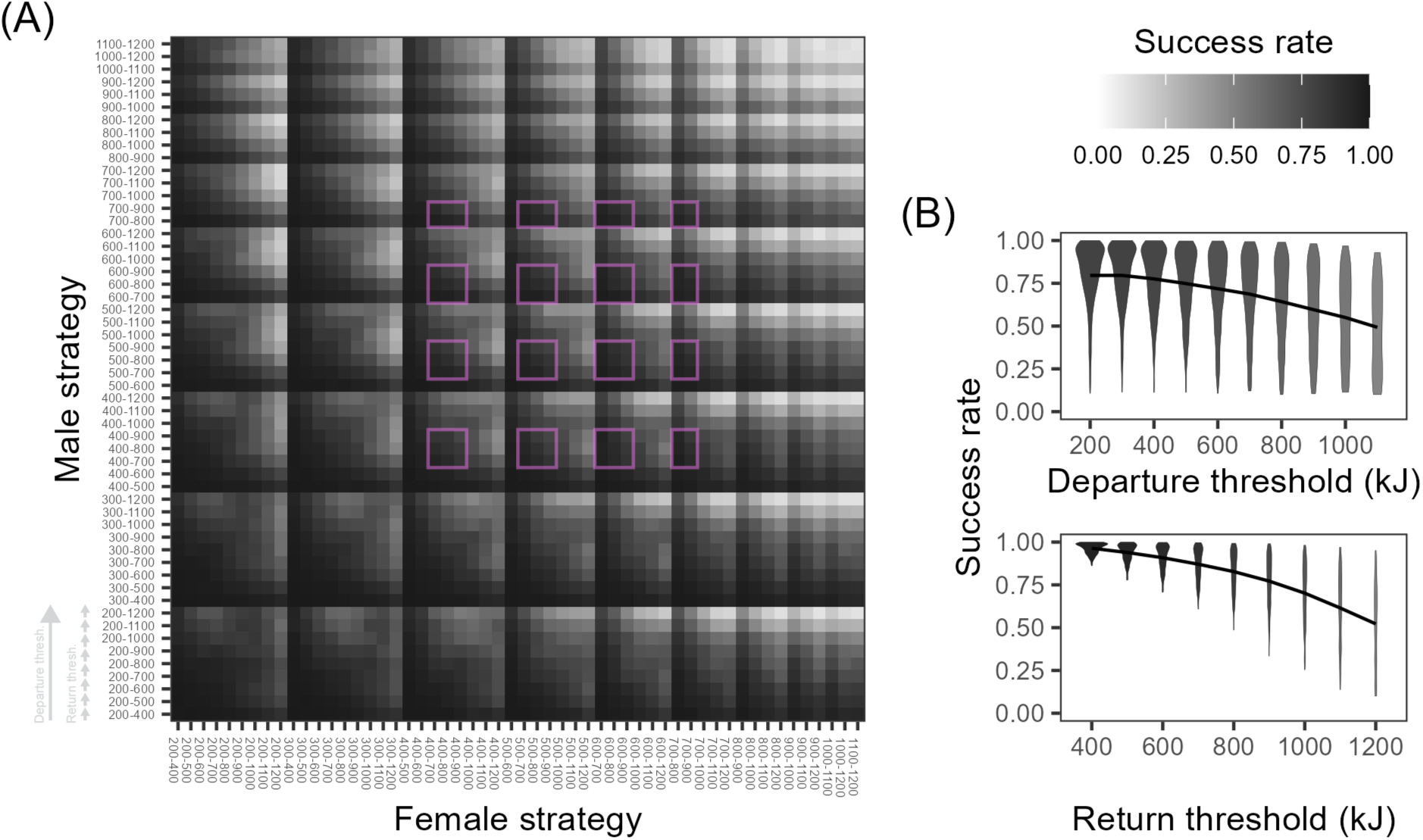
Parent strategies determine hatching success. Strategies are defined by the departure and return thresholds that parents use to switch between incubating and foraging. (A) Success rates calculated from 1,000 iterations per pair strategy combination given empirical environmental parameters. Along the axes, parent strategies are ordered first by departure threshold then by return threshold (both ascending). Pink outlines indicate the empirical range of pair strategy combinations (400–700 kJ departure, 700–900 kJ return). (B) Generous strategies, with low departure or return thresholds (i.e., adults are willing to reach lower body condition), achieve the highest hatching success. Violins show distribution of success rates for individual parameters, sampled across all other parent and partner threshold parameters, with black curve indicating mean.

In general, “generous” parents that maintained low energy levels were more successful than “selfish” parents that maintained large energy surpluses (Fig. 3A). Yet some highly successful pair strategy combinations could maintain high average parent energy, while others suffered low energy (Fig. 3A). Highly successful strategies also tended to hatch eggs quickly (Fig. 3B). However, there was a tradeoff between these two metrics: successful strategies that maintained high parent energy levels also left eggs unattended more regularly, and therefore hatched eggs more slowly (Fig. 3C). A doubling in average parent energy from 500 kJ to 1,000 kJ postponed hatch dates from 43 ± 4 days to 50 ± 3 days. Empirical pair strategy combinations were intermediate for both parent energy and hatch date; across the full parameter search range, strategies that were more successful than the empirical average tended to offer faster hatching but lower parent energy (Fig. S6).

**Figure 3.**
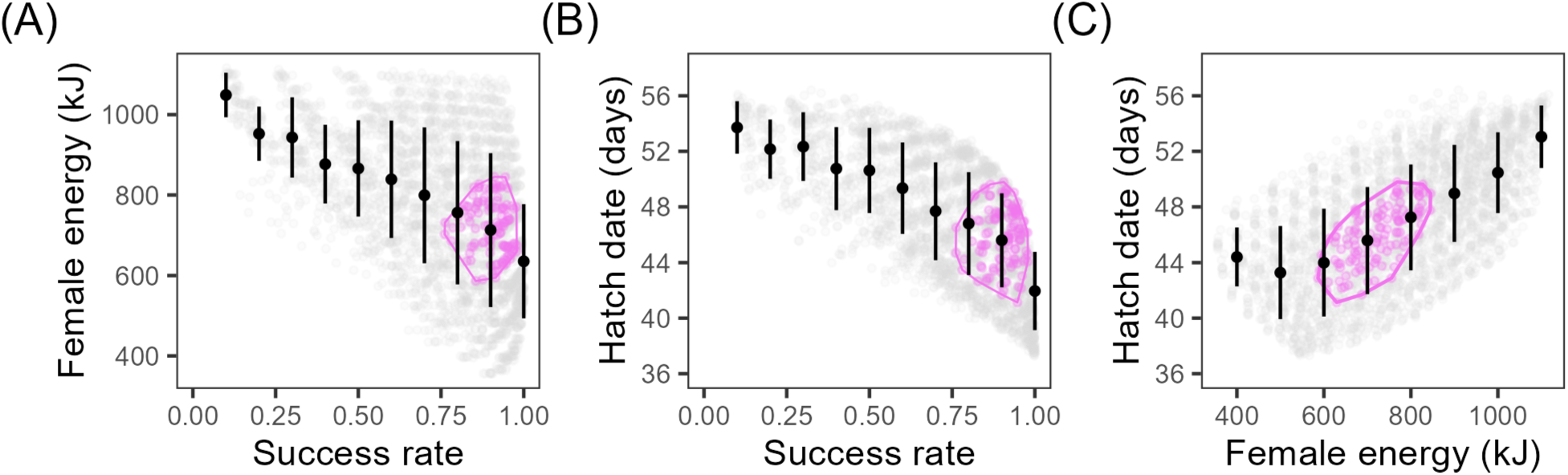
Parents navigate a tradeoff between their own energy and egg attendance. (A) In consistently successful strategies, parents maintain lower energy levels, but there is still high variation in energy levels for the most successful strategies. (B) Consistently successful strategies hatch eggs quickly. Because hatch date only increases when eggs are left unattended, higher hatch dates correspond to increased rates of egg neglect across the incubation period. (C) Strategies that allow parents to hatch eggs quickly require parents to maintain lower energy levels. In all panels, gray points show strategy combinations, defined by the departure and return thresholds of both parents. Pink points show empirical pair strategy combinations. Black points and lines show mean ± SD for bins across the x-axis. Energy and hatch date metrics are summarized for successful incubation only.

### Environmental conditions

There was a sharp drop in hatching success as environmental conditions declined, modeled as a reduction in the mean daily intake for foraging parents (Fig. 4). Looking across both empirical and non-empirical pair strategy combinations, hatching success dropped from 90.1 ± 14.2% in the most productive environment (foraging mean = 170 kJ/day) to <0.1% success in the most degraded environment (foraging mean = 130 kJ/day). A logistic regression indicated a switch-point (50% success rate) at 157 kJ/day.

**Figure 4.**
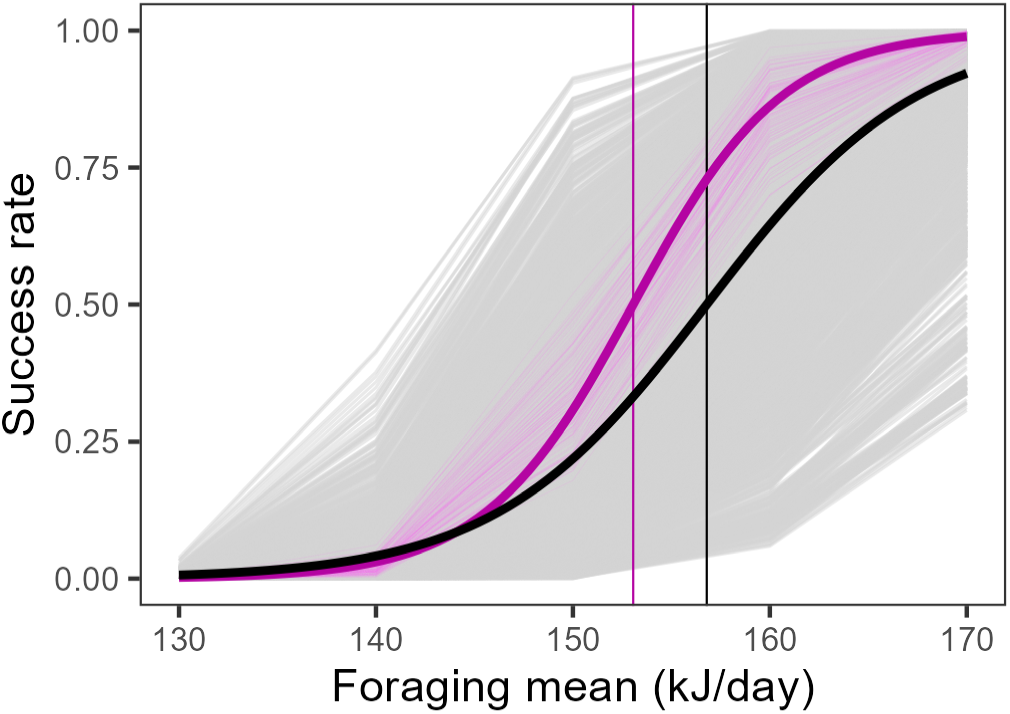
Environmental conditions, modeled as mean daily foraging intake, determine hatching success. Gray lines show results for each pair strategy combination, with empirical strategies in pink. Bold curves show logistic fit with vertical lines marking 50% success.

Empirical parent strategies were more successful than the full test range, with 98.8 ± 1.2% success in the 170 kJ/day environment, but still declined to <0.1% success at 130 kJ/day with a switch-point at 153 kJ/day (Fig. 4). Although empirical strategies approached perfect success in productive environments, additional simulations showed that success still required effort from both parents. Simulations with just one parent attempting to incubate resulted in 0% success until environmental conditions surpassed 200 kJ/day, with a switch-point at 260 kJ/day (∼160% of the empirical foraging intake values; Fig. S7).

The model identified cold shock as the major mechanism of incubation failure (Fig. 5A). In the most productive environments (foraging mean = 170 kJ/day), the average rate of cold shock failure was 9.6 ± 14.0%, which was more frequent than failure by slow development (0.3 ± 0.7%) or parent death (0%). As conditions worsened, the rate of cold shock increased rapidly to become the majority outcome (64.9 ± 23.0%) in the 150 kJ/day environment and the dominant outcome (97.5 ± 3.3%) in the 130 kJ/day environment. In contrast, the rate of failure from slow development peaked at 10.7 ± 12.0% in the 150 kJ/day environment. Parent death was rare, peaking at only 2.1 ± 3.1% in the 130 kJ/day environment.

**Figure 5.**
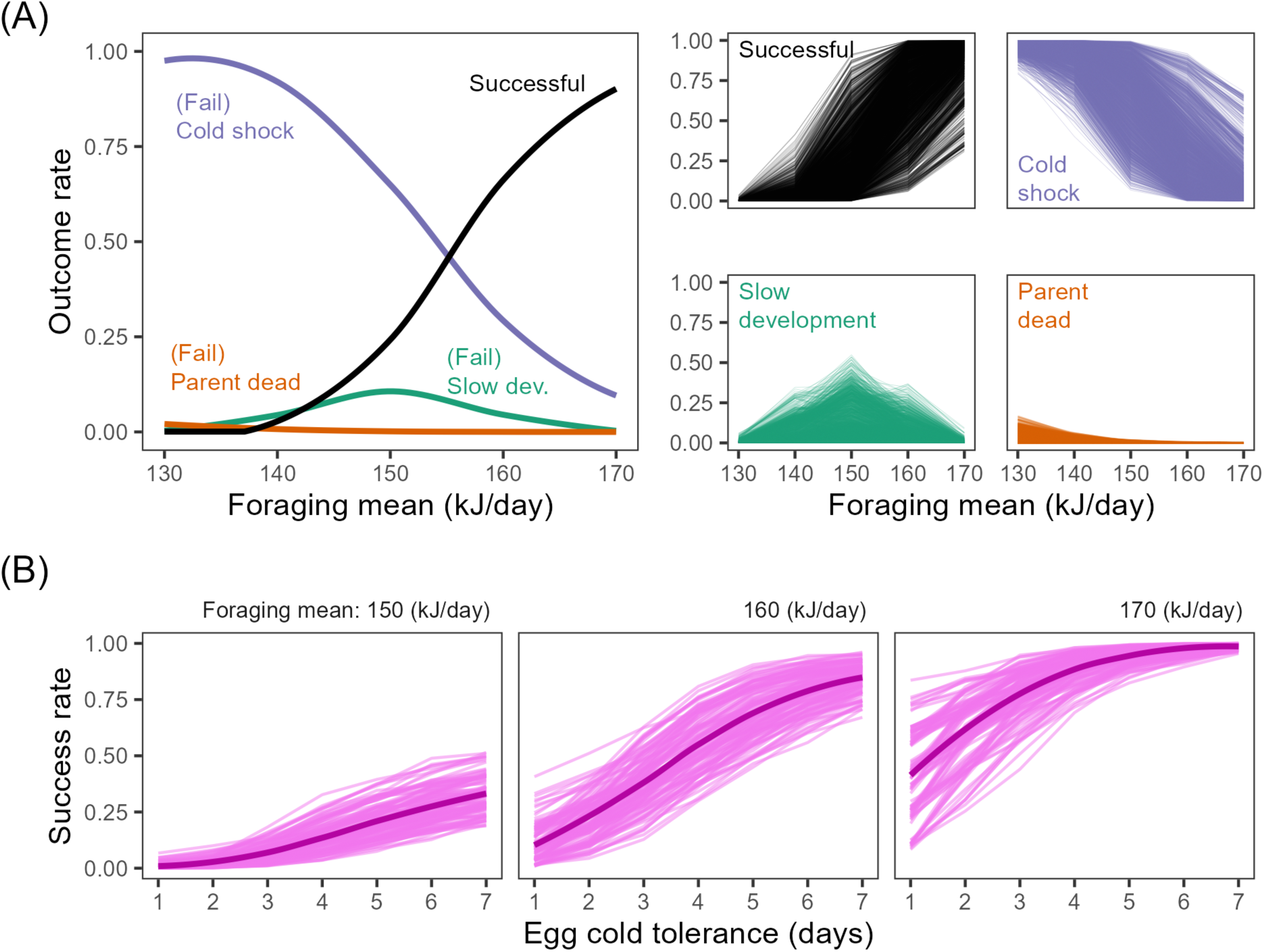
Cold shock is the primary point of incubation failure, making resilient eggs an essential factor in reproductive success. (A) As the environment degrades, incubation begins to fail due to a single point in the season when eggs are left unattended for too many days in a row (“cold shock”). Outcome rates summarized with locally weighted smoothing, with simulation results for each pair strategy combination on the right. (B) Even in productive environments, hatching success declines sharply if eggs are not cold tolerant. Cold tolerance defines the maximum number of consecutive days an egg can survive unattended before failing from cold shock. Pink lines show results for each empirical pair strategy combination, with curves from locally weighted smoothing.

The rate of cold shock failure in the base model is equivalent to the probability of a stretch of eight consecutive days of neglect occurring at least once during incubation (i.e., exceeding the egg cold tolerance reported in the wild; Table S1). Hatching success was thus highly sensitive to egg cold tolerance (Fig. 5B). Even in the most productive environment (foraging mean = 170 kJ/day), success rates for empirical pair strategy combinations dropped from 98.8 ± 1.0% when egg cold tolerance was seven days to just 41.4 ± 18.4% when the egg could withstand just one day unattended. Near empirical conditions (foraging mean = 160 kJ/day), hatching success was only 10.8 ± 7.9% given an egg cold tolerance of one day.

## DISCUSSION

Our first key finding is that simple energetic rules are largely sufficient to model the incubation schedules of Leach’s Storm-Petrels, an imperiled seabird with an energetically, spatially, and temporally extreme form of biparental care. Combining straightforward behavioral rules with empirical energetic parameters generates natural incubation rhythms (∼4**-**day bouts) and the high hatching success (∼90%) of some wild populations (Table 1). Experiments have found that the incubation schedules of birds are not determined by *physiological limits*: wild birds rarely end incubation bouts because they run out of energy (Bulla *et al*. 2015; Kosztolányi *et al*. 2009). Yet our results suggest that simple *energetic decisions* may still describe many facets of biparental care (Chaurand & Weimerskirch 1994; Gillies *et al*. 2022; Ricklefs 1983). A pertinent extension involves reparameterizing the model to test whether, or where, simple energetic scaling can explain the astonishing diversity of biparental care schedules across birds, mammals, fish, frogs, and even some insects (Bulla *et al*. 2016; Perrone & Zaret 1979; Townsend *et al*. 1984; Trumbo 2012; Woodroffe & Vincent 1994).

The limitations of our model are equally informative. We simulated a wide range of environments and parent strategies to understand the underlying structure of a biparental care system. This contrasts with a classic game-theoretic approach, which often frames biparental care as a dynamic negotiation between partners (McNamara *et al*. 2000). Our results suggest that seabirds do not need dynamic strategies to successfully hatch an egg. However, parents in the model leave the egg unattended at artificially high frequencies, and therefore take longer to hatch their eggs (Table 1). Thus, an important question is whether more complicated strategies can increase nest attendance, parent energy reserves, or both. Game theory raises a wide range of hypothetical strategies for dynamic partner coordination, retaliation, or compensation (Leimar & McNamara 2023; Royle *et al*. 2016). In seabirds, dynamic strategies may include compensating for reduced partner effort (Gillies *et al*. 2022), alternating between short- and long-distance foraging routes (Tyson *et al*. 2017; Wojczulanis-Jakubas *et al*. 2018), and balancing effort based on previous experience with the same partner (Patrick *et al*. 2020; van de Pol *et al*. 2006).

Another critical missing strategy was nest abandonment. Parents in the model could only breed or die. In reality, parents also have the option to abandon their young (McNamara *et al*. 2000). Indeed, nest abandonment is often a proximate cause of breeding failure in storm-petrels (Zangmeister *et al*. 2009). Our simulations indicate that a single storm-petrel cannot successfully incubate under realistic energetic conditions (Fig. S7). Thus, nest abandonment in this system is a true surrender of a reproductive opportunity rather than a strategic attempt to leave a partner holding the bag (Trivers 1972).

More complicated strategies may also be needed to explain the full range of sexual differences in biparental care. Our simulations recover a subtle sexual bias in the behavior of an otherwise monomorphic species, with males showing longer incubation bouts (Mauck *et al*. 2011). In the model, this bias emerges from the energetics and timing of egg-laying (Pinet *et al*. 2012; Tyson *et al*. 2022) rather than an evolutionary history of sexual conflict, mate choice, or sex-specific adaptation (Andersson 1994). However, the incubation bias in the model (0.2 days) is even smaller than empirical estimates (0.5 days). Larger biases towards male incubation would require an egg that is much more energetically or nutritionally costly than currently known (Fig. S3; Wojczulanis-Jakubas *et al*. 2014). Further, although we assumed that females start the season incubating because they must lay the egg, male Leach’s Storm-Petrels are often the ones actually found incubating first (S. Neirink, pers. comm.; cf. Pinet *et al*. 2012). Swapping the starting order in the model can reverse the automatic male bias in incubation effort (Fig. S4). Thus, some core aspects of behavior in this system cannot be explained by direct energetic flux alone and require wider evolutionary explanations.

Our second key finding is that generous parent strategies, with low departure or return thresholds, are the most successful. However, parents must navigate two levels of tradeoffs. Low departure or return thresholds offer higher hatching success but, by definition, run the risk of smaller energy reserves for parents (Figs. 2, 3A). Further, successful parents must still choose between maintaining high energy and hatching an egg quickly (Fig. 3). This secondary tradeoff appears because cold, unattended eggs have slower development (Boersma & Wheelwright 1979; Elliott *et al*. 2021) and generous strategies are more likely to attend the egg.

For parents, maintaining good condition may be important for survival to future breeding attempts and withstanding the other energetically intense phases of seabird reproduction—chick brooding and rearing—later in the season (Cruz-Flores *et al*. 2021; Erikstad *et al*. 1998; Ricklefs 1983). At the same time, earlier hatch dates may benefit chicks through phenological relationships with environmental resources, competitors, or predators (Arnold *et al*. 2004; Hipfner *et al*. 2010; Ramirez *et al*. 2016; but see Price *et al*. 1988). We note that the empirical parent strategies tested here for Leach’s Storm-Petrels were not the most successful across the parameter search range (Fig. S5). Instead, empirical strategies traded lower hatching success, along with a delay in hatch date, for elevated parent energy (Fig. S6). These results are consistent with the hypothesis that long-lived organisms such as storm-petrels weigh future reproductive opportunities more heavily than any individual breeding attempt (Mauck & Grubb 1995). Of course, real organisms may also behave suboptimally due to incomplete environmental information, limited physiological self-assessment, or evolutionary constraints on individually optimizing behavioral components of parental care (Pierce & Ollason 1987).

Our third key finding is that all parent strategies, generous or otherwise, collapse as the marine environment degrades. A decline in environmental conditions of ∼5% (from 162 to 153 kJ/day of mean foraging intake) drops the hatching success of empirical strategies from >90% to <50% (Fig. 4). At a ∼20% decline (to 130 kJ/day), the model suggests it is not possible for storm-petrels to successfully hatch an egg. These results help explain the ties between long-term declines in hatching success and rising global temperatures (Mauck *et al*. 2018). The dramatic effects of environmental degradation on reproductive success depend on our modeling assumptions, including the fact that parents are restricted to simple, static energetic decisions.

However, the results raise alarms about indirect causes of reproductive failure in imperiled seabirds. Like nearly all marine vertebrates, seabirds are in decline (McCauley *et al*. 2015). Many of the pressures facing seabird populations are direct threats, including introduced predators that eat chicks at nesting colonies and fishing lines that ensnare adults at sea (Dias *et al*. 2019). But recent work has emphasized more nuanced threats, such as nesting failures that cause seabirds to break up their long-term pair-bonds (Pollet *et al*. 2026; Sun *et al*. 2024; Ventura *et al*. 2021). The degradation of the marine environment may have indirect and chronic, as well as direct and acute, impacts on vulnerable populations.

Indeed, our fourth key finding is that reproductive failure in this system does not emerge directly, as through parent starvation, but rather indirectly through a “schedule breakdown.” As the environment degrades, it becomes increasingly likely that both parents spend too long foraging at the same time. The egg then dies of cold shock due to prolonged neglect (Fig. 5A). This failure can arise from a single period of miscoordination when parents are otherwise healthy and frequently attending the nest. Empirical work shows how environmental stressors, such as dehydration and heat stress, can reduce attendance and feeding rates (Levillain *et al*. 2025). Here, environmental deficits kill the offspring through a single scheduling error.

Studies of biparental care are generally focused on parent behavior, but the threat of cold shock draws our attention to the offspring. Simulations show that reproductive success plummets when eggs are less cold tolerant (Fig. 5B). In other words, the model predicts strong selection for resilient eggs. Storm-petrel eggs can withstand up to a week of neglect (Boersma & Wheelwright 1979; Jouventin *et al*. 1985). Most other species’ eggs are much more sensitive in the middle of incubation (Ahmad & Li 2023), although prolonged cold tolerance has convergently evolved in at least one other lineage of seabirds (*Synthliboramphus* murrelets; Murray *et al*. 1980). Our results suggest that cold-tolerant eggs are a fundamental adaptation in these lineages, allowing parents to pursue wide-ranging and uncertain foraging opportunities despite the risk of leaving the nest unattended. Future research is needed to understand the genetic, developmental, and physiological mechanisms that make some seabird embryos resilient to extreme conditions.

Overall, we emphasize the importance of investigating the patterns and mechanisms of reproductive schedules themselves (Bulla *et al*. 2016). As in classic life history theory, where fitness is revealed to unfold across a lifetime of development and senescence (Williams 1966), reproduction in a single season unfolds across a discrete schedule of events (Houston & McNamara 1999). A single event, such as a prolonged bout of neglect, can have consequences for parents and offspring alike. The outstanding question is how organisms craft such effective and resilient schedules in such an uncertain world.

## ACKNOWLEDGMENTS

Mary Lou Zeeman, Elizabeth Marschall, and Alvaro Sanchez provided important feedback on earlier versions of the model. We are grateful to Richard Prum, Ingrid Pollet, and the students, faculty, and staff at the Bowdoin Scientific Station on Kent Island for discussions on the topic.

## AI DECLARATION

We used Claude (Opus/Sonnet 5) to verify the codebase, implement simulation results processing, and flag typographical, syntactical, and consistency errors in the manuscript.

## DATA AVAILABILITY

Model and analysis code is available on GitHub (https://github.com/ltaylor2/Petrel_Schedule).

## STATEMENT OF AUTHORSHIP

LT and RM conceived of the study. LT developed and implemented the model and analysis with input from PJ, MH, and RM. LT wrote the first draft of the manuscript, and all authors contributed substantially to revisions.

## Notes

https://github.com/ltaylor2/Petrel_Schedule

### Competing Interest Statement

The authors have declared no competing interest.

