## Supplementary Material for "Scheduling problems and the energetics of biparental care in a model of imperiled seabirds"

**TABLE OF CONTENTS**


p. 1 Table S1
p. 2 Figure S1
p. 3 Figure S2
p. 4 Figure S3
p. 5 Figure S4
p. 6 Figure S5
p. 7 Figure S6
p. 8 Figure S7
p. 9 References for supplementary material

**Table S1.** Energetic parameters for Leach's Storm-Petrels in the Northwest Atlantic

| **Parameter** | | **Empirical value** | **Test range** |
|---|---|---|---|
| *Parent* | Basal metabolic rate | 52 kJ/day[A,B] | |
| | Active metabolic rate | 123 kJ/day[A] | |
| | Departure threshold | 400–700 kJ[A] | 200–1,100 kJ |
| | Return threshold | 700–900 kJ[A] | 400–1,200 kJ |
| | Season starting energy | 766 kJ[C] | 300–1,300 kJ |
| *Egg* | Egg cost to females | 69.7 kJ[D] | 0–500 kJ |
| | Starting hatch date | 37 days[E] | |
| | Hatch date limit | 60 days[F] | |
| | Egg cold delay | 1.43 days/day[G] | |
| | Egg cold tolerance | 7 days[F,G] | 1–7 days |
| *Environment* | Foraging condition mean | 162 kJ/day[H] | 130–170 kJ/day |
| | Foraging condition SD | 47 kJ/day[H] | |

[A]Ricklefs *et al.* (1986); [B]Blackmer *et al.* (2005); [C]Taken as average starting energy at the beginning of an incubation bout, Ricklefs *et al.* (1986); [D]Montevecchi *et al.* (1983); [E]Pollet *et al.* (2021); [F]*Unpublished data*, Kent Island, New Brunswick, Canada; [G]From Fork-tailed Storm-Petrel (*Hydrobates furcatus*), Boersma & Wheelwright (1979); [H]Montevecchi *et al.* (1992).

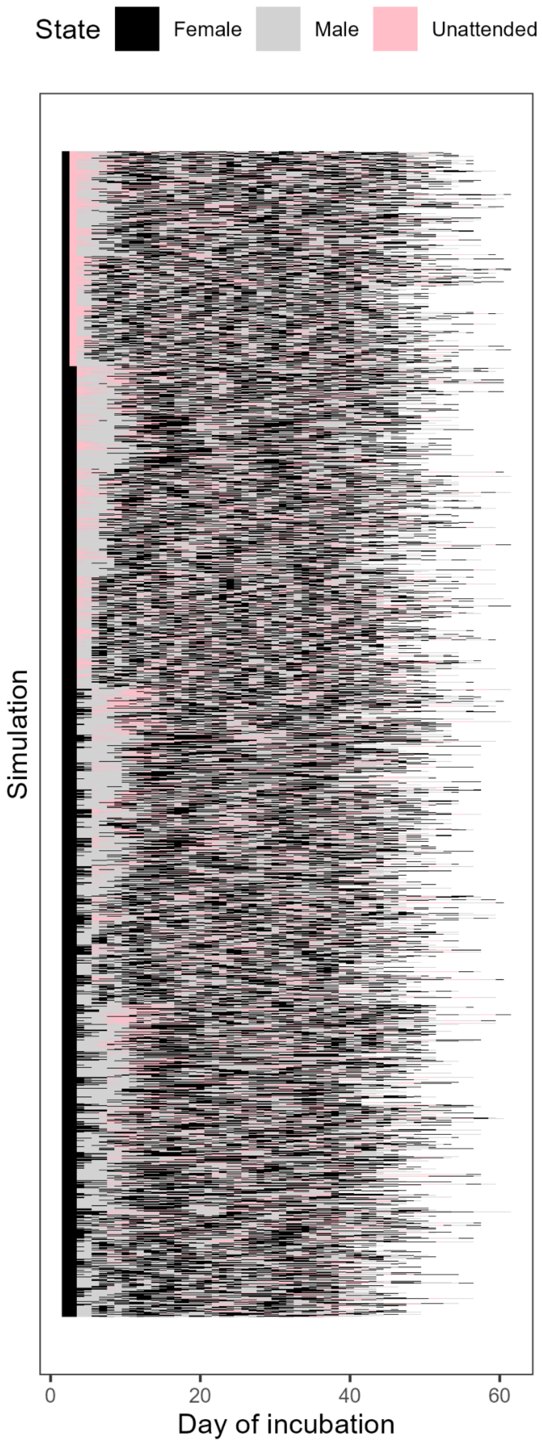


**Figure S1.** Simulated incubation schedules for Leach's Storm-Petrels given empirical parameters for the environment, egg, and the range of parent strategies. Runs are ordered by pair strategy combination. Only successful schedules are shown, meaning all simulations end at hatch.

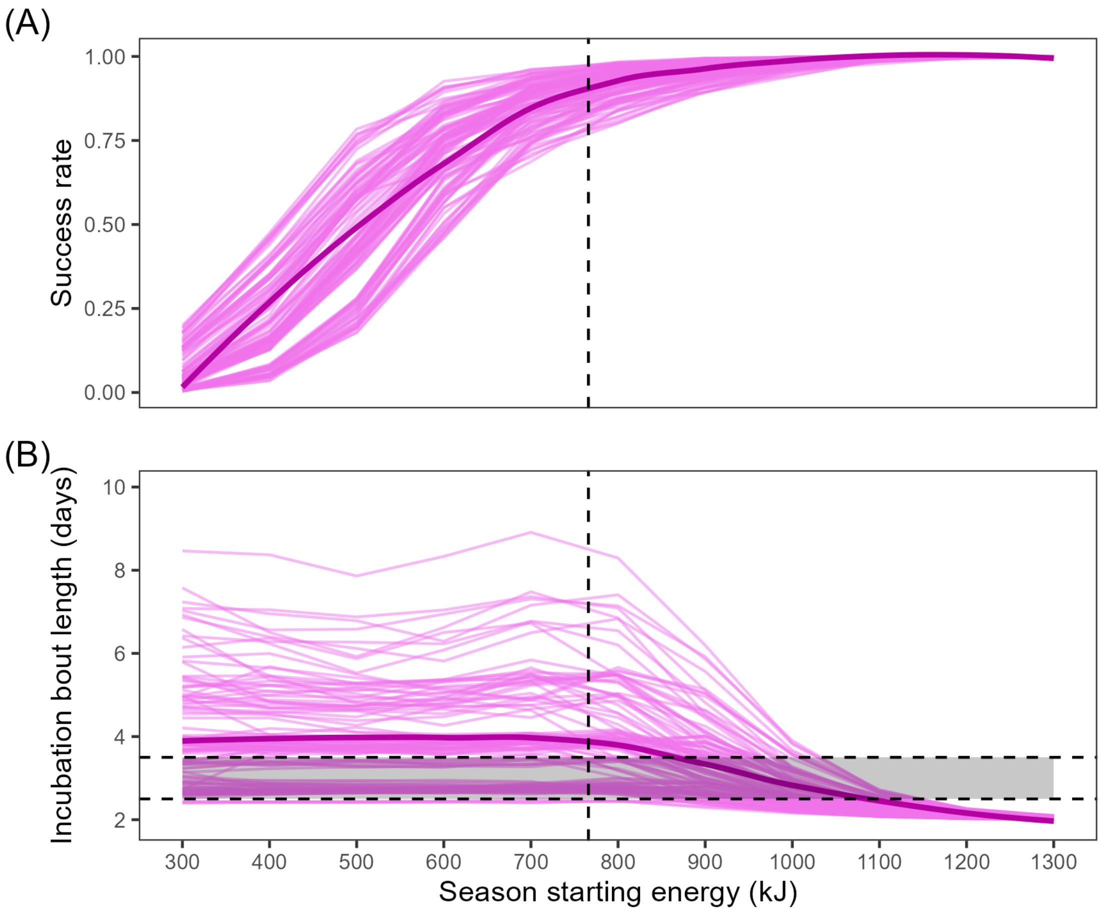


**Figure S2.** Season starting energy influences hatching success and incubation rhythms during simulated incubation. Simulations run across a range of season starting energy levels (300–1,300 kJ, empirical estimate = 766 kJ) using empirical parameters for the environment, egg, and the range of parent strategies. (A) Increasing season starting energy raises overall hatching success. Pink lines show results from each empirical pair strategy combination, with local smoothing in bold. (B) Starting with more energy reduces the average length of incubation bouts across the season. Dashed lines indicate empirical values, with gray rectangle highlighting the range of average bout lengths for wild populations.

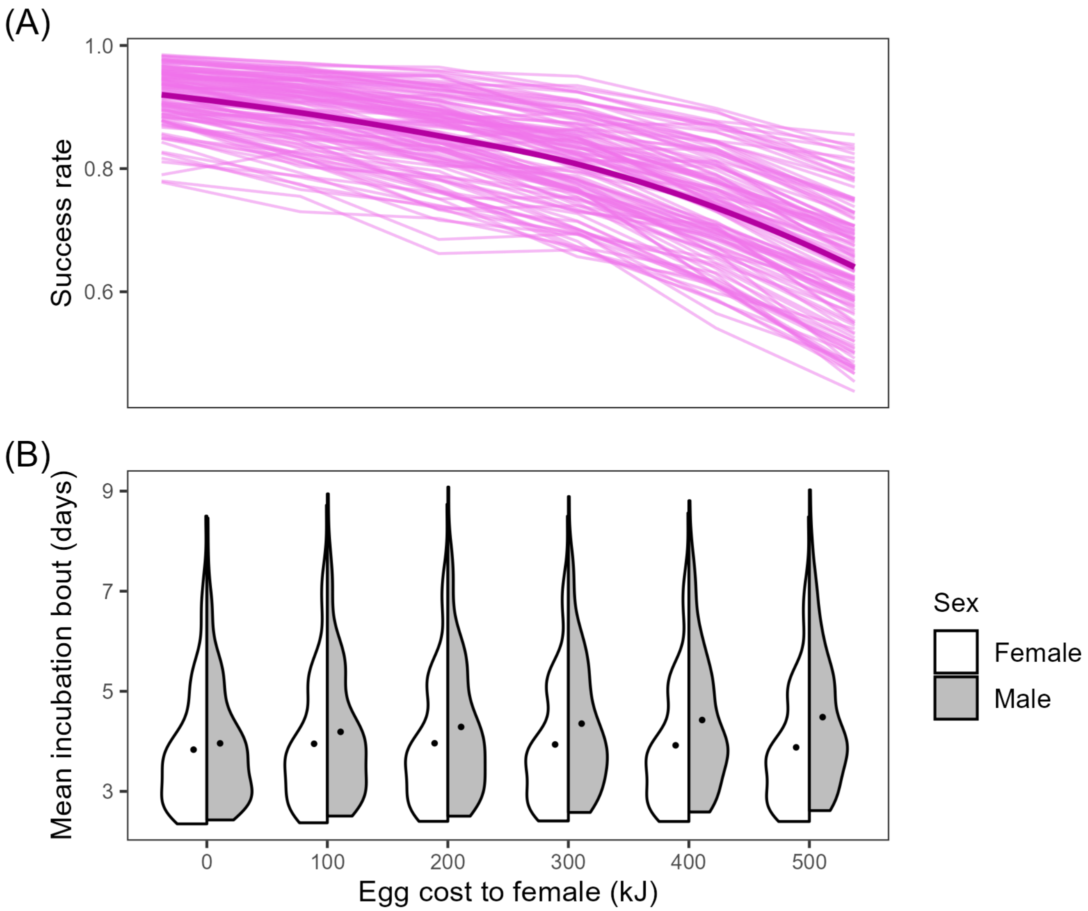


**Figure S3.** The energetic cost of the egg to females contributes to a male bias in incubation attendance. Simulations run across a range of egg costs (0–500 kJ, empirical estimate = 69.7 kJ) using empirical parameters for the environment and the range of parent strategies. (A) Increasing egg costs results in lower hatching success. Pink lines show results from each empirical pair strategy combination, with local smoothing in bold. (B) More expensive eggs result in a greater male bias during incubation. Violins show the distribution of mean incubation bout lengths for males and females, excluding the first and last bout for each parent, across all empirical pair strategies.

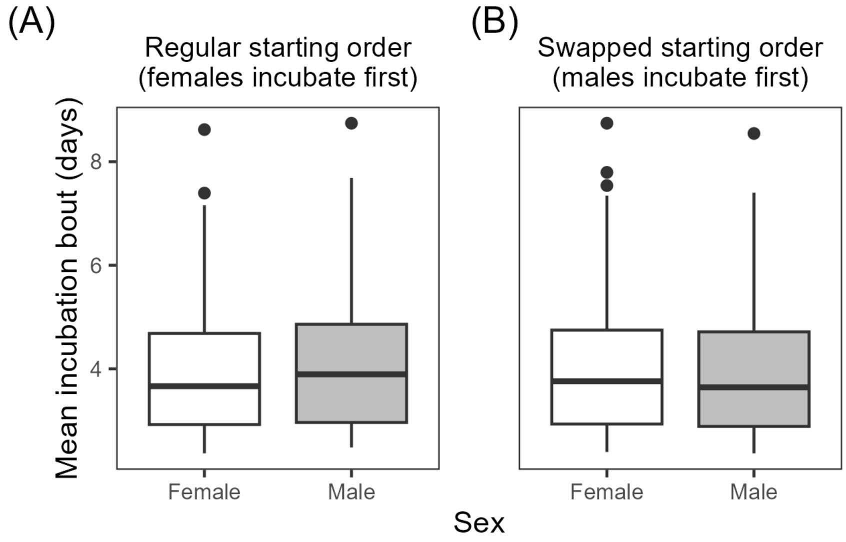


**Figure S4.** Initial state assignments for males and females contribute to a male bias in incubation attendance. Simulations used empirical parameters for the environment, egg, and the range of parent strategies. Summary values exclude the first and last bout for each parent. (A) In the regular model, where females begin the season incubating and males begin foraging, males average slightly longer incubation bouts (4.1 ± 1.3 days) than females (3.9 ± 1.3 days). (B) The bias reverses with smaller magnitude when initial states are swapped (males: 3.9 ± 1.3 days, females: 4.0 ± 1.3 days). In both panels, central lines indicate median, boxes give 25–75th percentiles, whiskers give 1.5 times inter-quartile range, and points indicate outliers.

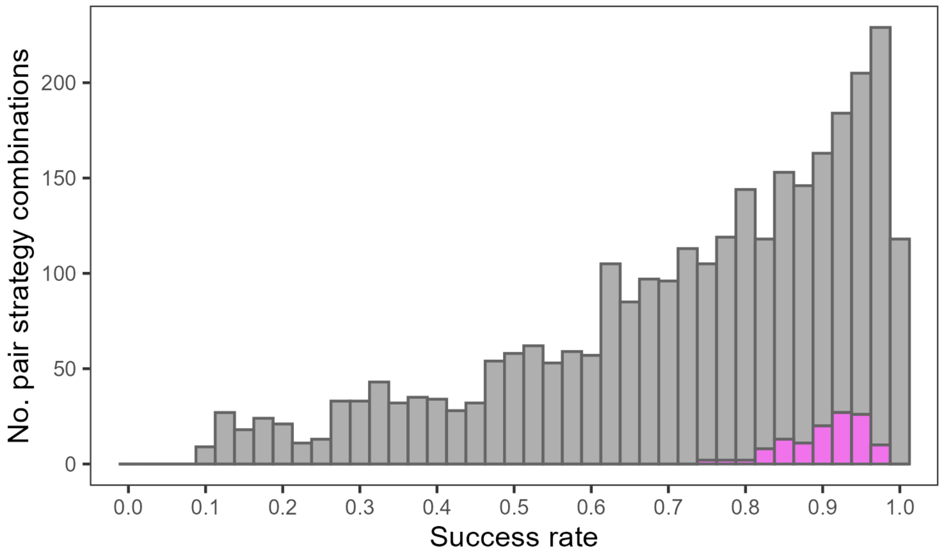


**Figure S5.** Hatching success for pair strategy combinations given empirical parameters for the egg and environment. Pink indicates empirical strategies. Bars are stacked.

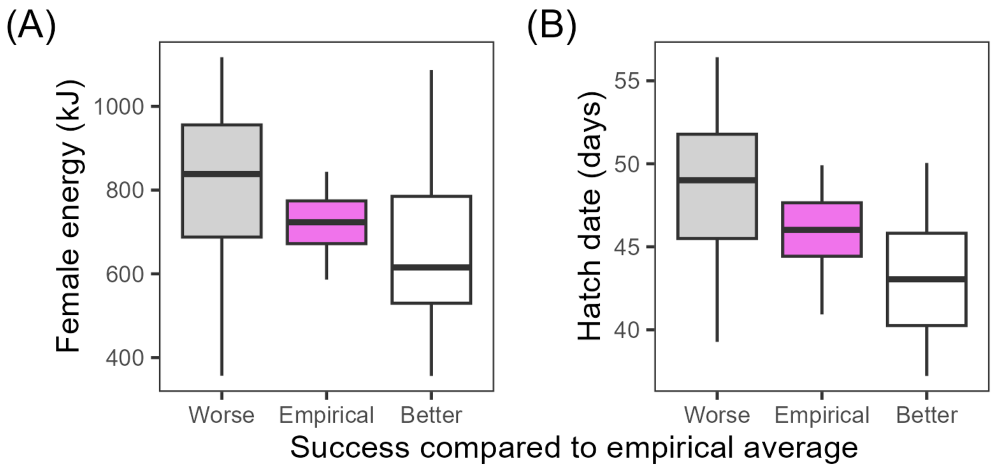


**Figure S6.** Empirical pair strategy combinations are intermediate for the secondary metrics of parent energy and hatch date. Boxes show pair strategy combinations split into three categories: empirical strategies, non-empirical strategies with a success rate that is better than the average of empirical strategies, and non-empirical strategies with a success rate that is worse than the average of empirical strategies. (A) Parent energy, summarized as mean female energy throughout the incubation period. (B) Hatch date, which is determined by the attendance rate of the egg. In both panels, central lines indicate median, boxes give 25–75th percentiles, and whiskers give 1.5 times inter-quartile range.

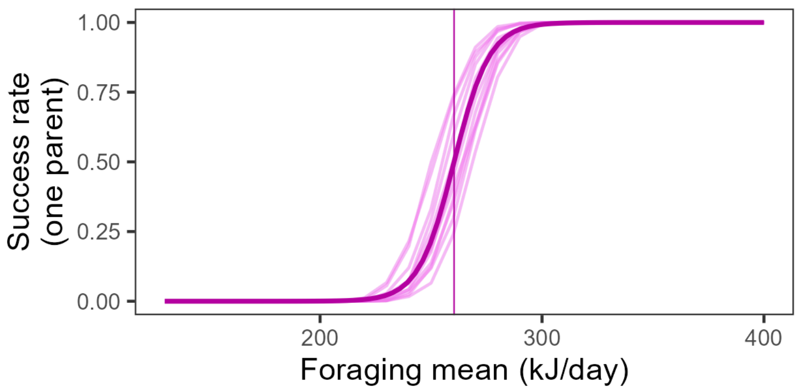


**Figure S7.** One storm-petrel cannot hatch an egg under realistic environmental conditions. Pink lines show results from each empirical parent strategy given only one parent (female), with bold curve giving the logistic fit and vertical line marking 50% success. All other parameters were held at empirical estimates.
